# Plasmids for independently tunable, low-noise expression of two genes

**DOI:** 10.1101/515940

**Authors:** João P. N. Silva, Soraia Vidigal Lopes, Diogo J. Grilo, Zach Hensel

**Author notes:** Funded by Project LISBOA-01-0145-FEDER-007660 (Microbiologia Molecular, Estrutural e Celular) (Zach Hensel).

## Abstract

Some microbiology experiments and biotechnology applications can be improved if it is possible to tune the expression of two different genes at the same time with cell-to-cell variation at or below the level of genes constitutively expressed from the chromosome (the “extrinsic noise limit”). This was recently achieved for a single gene by exploiting negative autoregulation by the tetracycline repressor (TetR) and bicistronic gene expression to reduce gene expression noise. We report new plasmids that use the same principles to achieve simultaneous, low-noise expression for two genes in *Escherichia coli*. The TetR system was moved to a compatible plasmid backbone, and a system based on the *lac* repressor (LacI) was found to also exhibit gene expression noise below the extrinsic noise limit. We characterized gene expression mean and noise across the range of induction levels for these plasmids, applied the LacI system to tune expression for single-molecule mRNA detection in two different growth conditions, and showed that two plasmids can be co-transformed to independently tune expression of two different genes.

## Introduction

We recently reported the development of a plasmid-based gene-expression system in which a gene of interest was expressed bicistronically with the tetracycline repressor (TetR) [1]. Using this gene expression system, cell-to-cell variation was below the “extrinsic noise limit” (coefficient of variation squared of protein concentration, *CV^2^* ≈ 0.1) observed for genes expressed from the chromosome [2]. When TetR and GFP were expressed bicistronically, GFP induction and gene expression noise were similar to those observed for a TetR-GFP fusion protein with autoregulation [3]. Compared to induction of gene expression under the control of a constitutively expressed transcriptional repressor, the inducer dose-response was relatively linearized, and gene expression noise was much lower at intermediate induction levels [1].

Our experiments in mRNA detection and other single-molecule applications in living *E. coli* cells sometimes require the tunable expression of two different genes, both with low noise levels. For example, adopting a recently reported mRNA detection system based on local enrichment of fluorescent RNA-binding proteins [4] for use in *E. coli* requires lower noise in protein production relative to the same system in *S. cerevisiae* because of a smaller cell volume for *E. coli* and the inability to sequester unbound protein in the nucleus. At the same time, tunable expression with low noise in the level of the target RNA is desired to make it possible to characterize the accuracy of RNA detection over a range of RNA levels. We hoped that expressing both the target RNA and RNA-binding fluorescent protein on two plasmids that could be tuned independently would simplify and accelerate development of new RNA-detection systems in *E. coli.* Achieving this was a three-step process: first, characterizing the TetR-based system on a compatible plasmid backbone; second, establishing an orthogonal, low-noise expression system based on the *lac* repressor (LacI); and third, showing that the two systems can be tuned independently.

## Results

### Moving bicistronic autoregulatory construct to a compatible plasmid backbone

The first step in creating a low-noise system for tuning expression of two genes was to establish that a previously characterized, bicistronic autoregulatory circuit functions well in a compatible plasmid backbone. In this expression system, GFP and TetR are expressed bicistronically from the TetR-repressible promoter P_LtetO-1_ and expression is induced by the addition of ATc [1]. This system was shown to have low noise and a linearized dose response compared to a system in which TetR was constitutively expressed. We moved the system from a plasmid with a p15A replicon conferring ampicillin resistance to a lower-copy-number plasmid with a pSC101 replicon conferring spectinomycin resistance [5]. The p15A and pSC101 replicons have been used together in multiplasmid systems [6].

GFP expression mean and noise were characterized from low to high levels of induction by flow cytometry. Figure 1 shows that pJS101 induces at similar ATc concentrations as pZH509, with the change to the lower-copy pSC101 backbone resulting in a 58% drop in mean expression levels at a wide range of ATc concentrations. For a similar expression system in the absence of autoregulated TetR expression, moving the P_LtetO-1_ promoter from a p15A to a pSC101 backbone resulted in an 87% drop in expression [7]. A smaller change was expected in our experiment, since negative autoregulation provided dosage compensation, just as autoregulation can reduce noise in plasmid copy number [3,8,9].

**Figure 1:**
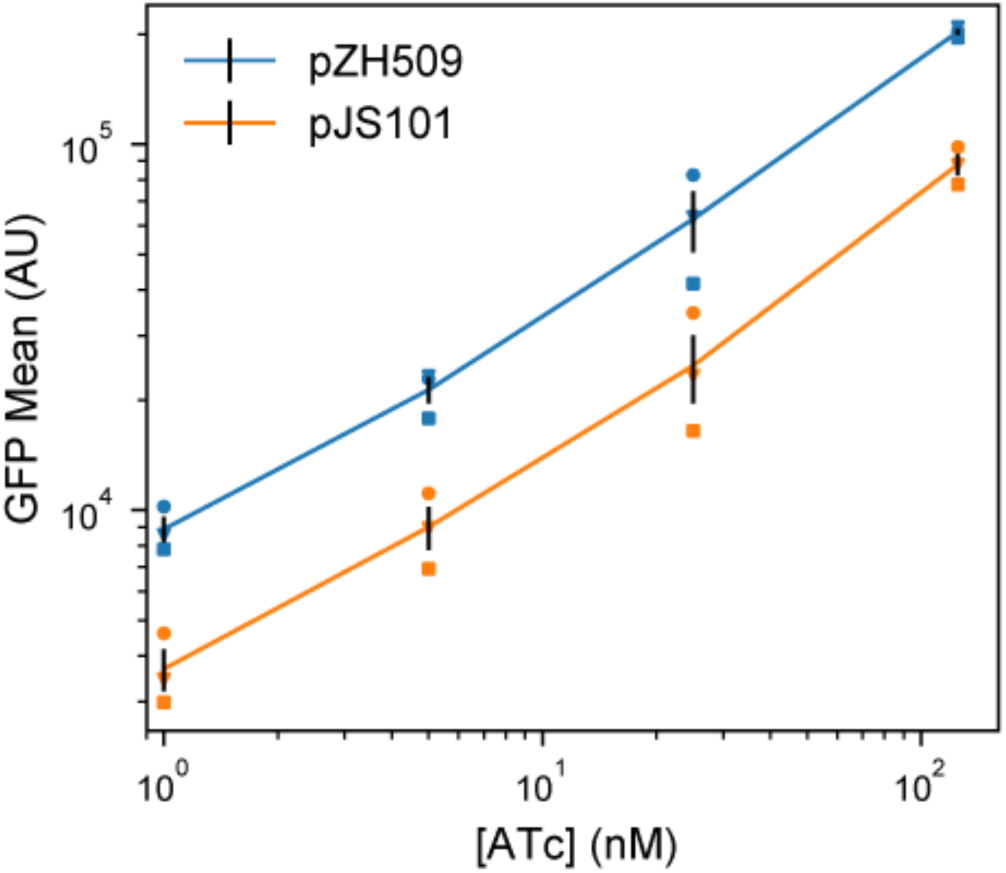
Moving the TetR expression system to a compatible plasmid backbone. Cultures of *E. coli* MG1655 harboring pZH509 (p15a origin, blue) or pJS101 (pSC101 origin, orange) expressing GFPmut2 with bicistronic autoregulation by TetR were grown at 30 °C in rich media with induction of 1, 5, 25 and 125 nM ATc. Mean single-cell GFP fluorescence was estimated using flow cytometry. Mean GFP levels in 3 independent replicates are indicated with different shapes. Black lines indicate the mean plus or minus 1 standard error of the mean from the 3 replicates.

### An alternative regulatory construct with LacI replacing TetR

We hypothesized that replacing P_LtetO-1_ with the IPTG-inducible promoter P_LlacO-1’_ with similar characteristics [7], and replacing TetR with LacI might result in a similarly useful expression system that could be tuned independently. However, regulatory parameters for TetR and LacI vary significantly. TetR binds *tetO2* more strongly than LacI binds *lacO1* (approximately 0.5—1.0 kcal/mol difference in binding energy [10,11] for a single site, with 2 tandem sites in our constructs). And, TetR binds ATc much more strongly than LacI binds IPTG (over 3 orders of magnitude difference in typical concentrations required for half-induction [12,13]).

We first characterized induction of GFP expression in MG1655 cells harboring IPTG-inducible pJS102 by flow cytometry. Figure 2a shows an induction range of almost 2 orders of magnitude from 0 to 1250 μM IPTG, with very good reproducibility of induction levels in 3 independent experiments. Previous experiments with the TetR-based system showed a similar total dynamic range, but with a large jump in expression going from 0 nM to 0.5 nM ATc [1]. This effect is not seen for pJS102, suggesting that switching from TetR:ATc to LacI:IPTG improves the dynamic range of tunable induction levels to a small extent.

**Figure 2:**
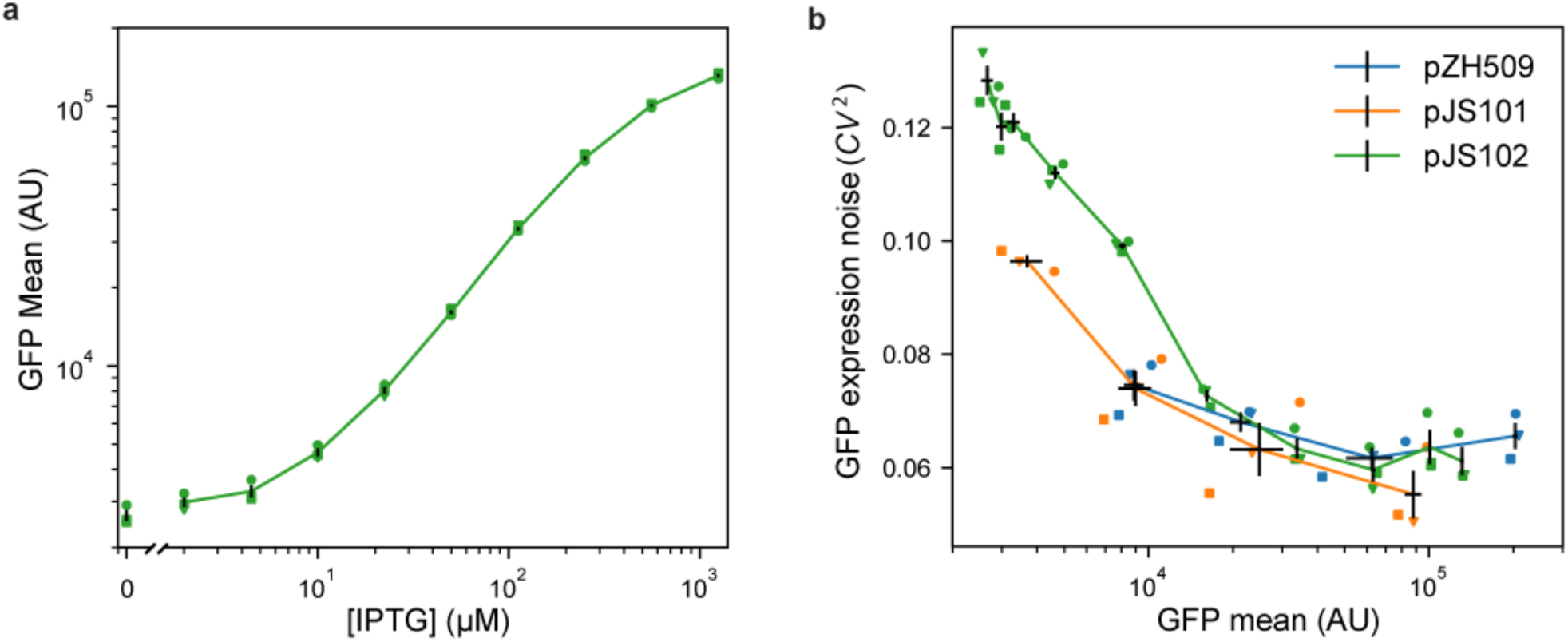
Characterizing mean expression levels and noise for different gene expression systems. Cultures of *E. coli* MG1655 harboring pJS102 (LacI regulation, p15a origin, green), pZH509 (TetR regulation, p15a origin, blue) or pJS101 (TetR regulation, pSC101 origin, orange) were grown at 30 °C in rich media with induction by IPTG (0, 2, 4.5, 10, 22.4, 50, 111.8, 250, 559 and 1250 μM) or ATc (1, 5, 25 and 125 nM). Mean single-cell GFP fluorescence and GFP expression noise was estimated using flow cytometry. Data from 3 independent replicates are indicated with different shapes. Black lines indicate the mean GFP expression and expression noise plus or minus 1 standard error of the mean from the 3 replicates. (**a**) Mean GFP expression for pJS102 at different IPTG concentrations shows tunable and reproducible induction over an ~50-fold dynamic range. Expression at zero IPTG is plotted separately to fit on logarithmic scale. (**b**) GFP expression noise as a function of mean for pZH509, pJS101 and pJS102. For all samples, GFP mean increases monotonically with inducer concentration. Note that the full range of induction is not shown here for pZH509 and pJS101 as even 1 nM ATc induces expression. GFP expression noise is low in all conditions for all strains.

Next, we compared noise in protein expression, with the concern that the *lac* operon present in the MG1655 host strain could lead to all-or-none expression at intermediate IPTG concentrations [14]. However, Figure 2b shows low noise in GFP expression at all IPTG concentrations, with noise levels comparable to pZH509 and pJS102 at the same mean GFP levels. Note that apparently high noise at very low expression was partially due to measurement noise, and, at any rate, was much lower than noise when expression is regulated by a constitutively expressed repressor [1].

We found that side scattering was weakly correlated with fluorescence, and thus with cell size, so gating for scattering modestly reduced measured noise in fluorescence intensity. However, we are comparing to an “extrinsic noise limit” determined from measurements of cell fluorescence divided by cell area [2], which effectively does the same thing. In practice, we observed slightly lower noise measurements for GFP concentration in fluorescence microscopy images compared to total GFP fluorescence in the gated flow cytometry sample for similarly induced strains. This difference was more significant for very-low-expression conditions, and noise in conditions where GFP fluorescence distributions significantly overlapped with ungated background events (GFP intensity less than 10^4^ in Figure S1) was somewhat overestimated. Our noise measurements were also consistent with the lower limit of gene expression noise found in many *E. coli* promoters using a similar flow cytometry method with similar gating and fitting procedures [15].

### Using the new induction system for detection of single mRNA in living *E. coli*

Recently, an improved method for detection of mRNA by local enrichment of fluorescent RNA-binding proteins was reported in *S. cerevisiae* [4]. This reduced the aggregation of mRNAs bound by the bacteriophage MS2 coat protein, which has also caused mRNA immortalization that has limited experiments in *E. coli* to observing transcription just after induction [16]. We hypothesized that aggregation could be reduced by reducing the expression levels of both mRNA and mRNA-binding proteins, and by having low cell-to-cell variation in expression. We developed a strain in which mRNA molecules encode mVenus-Cro and also include 24 tandem repeats of the binding sequence for the PP7 coat protein (PP7cp) [17]. These mRNAs were constitutively expressed at low levels (less than 1 molecule per cell). Plasmid pJS102 was used as a template to develop a fluorescent, IPTG-inducible reporter of expression, PP7cp-SYFP2.

We tested the utility of this expression system for tuning low-noise gene expression in different growth conditions. In previous experiments, we expressed the RNA-binding protein from a constitutive promoter integrated into the chromosome; this required long cycles of optimization every time a parameter was changed (e.g. growth media, temperature, and fluorescent protein sequence) that changed protein expression levels. Figure 3a shows that single-molecule mRNA detection was optimal (fluorescent mRNA spots are sufficiently bright but not obscured by background) at 100 μM IPTG in minimal media supplemented with 1% rich media. We note the absence of pole-localized fluorescent spots that characterize mRNA aggregation [18], and we observed reasonable mRNA lifetimes of a few minutes in time-lapse imaging.

**Figure 3:**
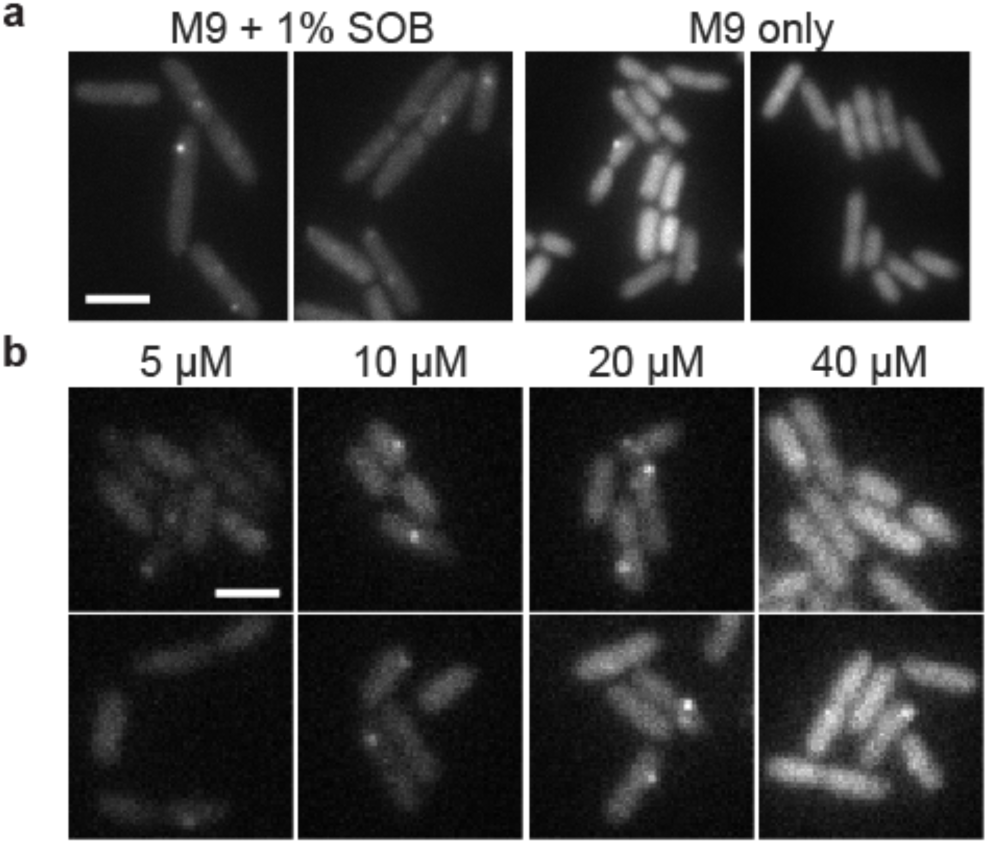
Using IPTG to tune expression of a fluorescent RNA-binding protein for single mRNA detection in different growth conditions. ZHX99 cells were grown with low expression levels (less than 1 molecule per cell) of mRNAs harboring 24 tandem binding sites for the PP7 coat protein fused to SYFP2. Cells were spotted on agarose gel pads and YFP fluorescence images were acquired. Two example images are shown for each condition. Images were taken shortly after preparing samples, so adjacent cells were not usually closely related in cell lineages. (**a**) PP7cp-SYFP2 was induced with 100 μM IPTG to detect single mVenus-Cro mRNA molecules in supplemented media and minimal media conditions; in minimal media there was too high a PP7cp-SYFP expression level to see single mRNA spots above background. Scale bar 4 μm. (**b**) Using the pJS102 expression system, PP7cp-SYFP2 expression levels were varied by induction with 5, 10, 20 and 40 μM IPTG. The range of 10—20 μM IPTG was identified to give bright mRNA spots above the background of unbound PP7cp-SYFP2 molecules. Scale bar 2 μm.

We moved to minimal media to explore a growth condition with different mRNA expression levels and slower growth rates, but found that 100 μM IPTG gave a background of unbound PP7cp-SYFP2 molecules that often made it impossible to detect mRNA spots. Figure 3b shows how the IPTG-inducible expression system made it simple to quickly scan different PP7cp-SYFP2 induction conditions and identify 10-20 μM IPTG as a range in which PP7cp-SYFP2 levels were high enough to label single mRNAs, but not so high as to give a high background of unbound molecules. Lastly, we note that the strain used for mRNA imaging has its entire *lac* operon replaced by the synthetic construct. Thus, this expression system works well both in the presence and absence of the *lac* operon.

### Independent, tunable expression of two genes

Lastly, we tested whether ATc-inducible and IPTG-inducible plasmids could be combined to achieve low-noise expression of two genes in the same cell. We replaced GFPmut2 in pJS102 with the fast-maturing RFP mScarlet-I [19] to create the plasmid pDG101. This plasmid was co-transformed with pJS101 into *E. coli* MG1655, and green and red fluorescence were compared at different combinations of ATc and IPTG concentrations. Figure 4a shows that pJS101 induction by ATc was unaffected by pDG101 induction by IPTG, and that all conditions gave low noise in GFP concentration. Figure 4b shows that mScarlet-I expression from pDG101 was similarly unaffected by the level of pJS101 induction by ATc. Figure 4c and 4d shows that mean expression levels were reproducible in 3 independent replicates, with low noise in each sample across a 5-fold increase in expression levels. There is some day-to-day variation in mean expression levels, but for each replicate induction of GFPmut2 and mScarlet-I is independent. Thus, independent, tunable expression of two genes can be achieved by replacing GFPmut2 in pJS101 and pJS102 with other genes of interest and co-transforming the plasmids into *E. coli.*

**Figure 4:**
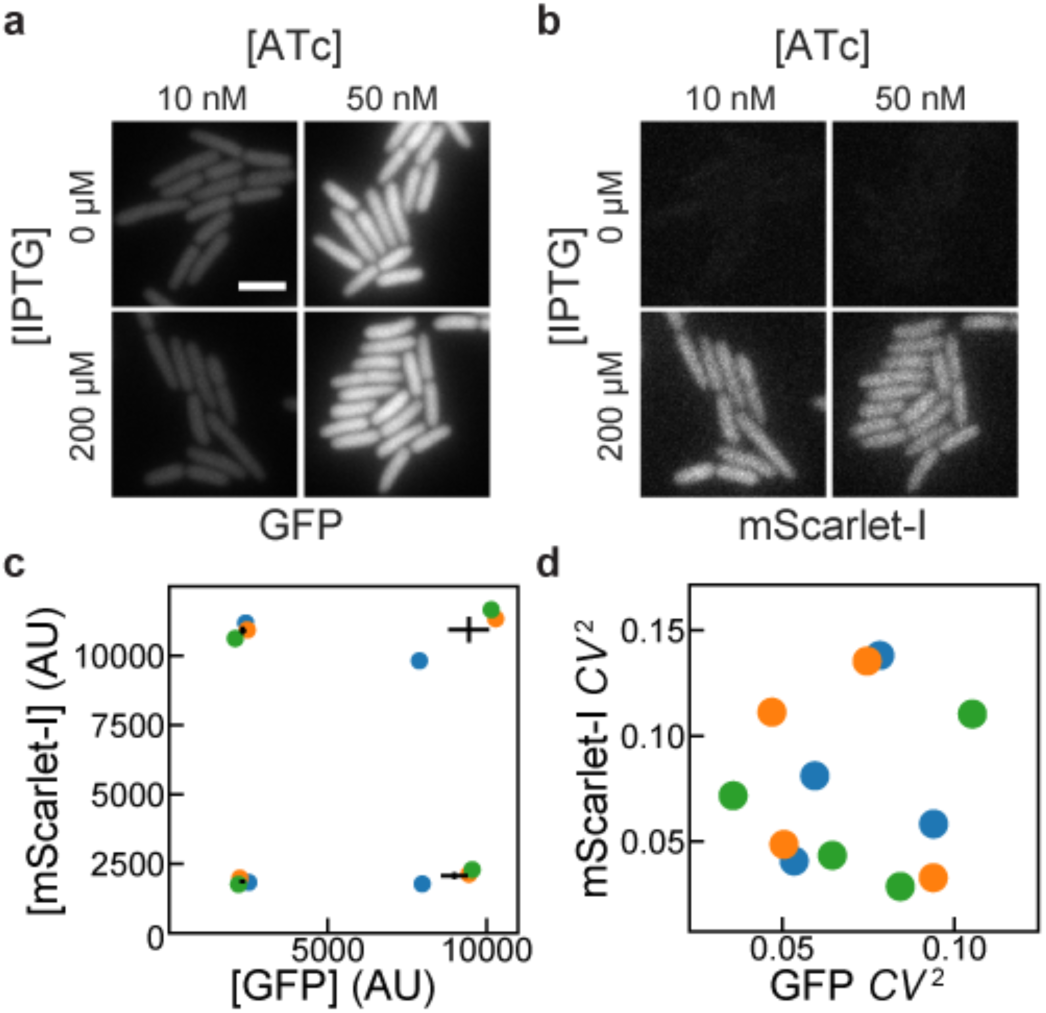
Independence of induction of TetR and LacI systems. MG1655 cells harboring pJS101 (ATc-inducible expression of GFP, pSC101 origin) and pDG101 (IPTG-inducible expression of mScarlet-I, p15a origin) were grown with different combinations of IPTG and ATc concentrations. GFP and mScarlet-I fluorescence was observed and quantified by fluorescence microscopy. (**a**) GFP fluorescence showed no apparent influence of IPTG on ATc-induced expression of GFP for cells grown at 30 °C in EZ-RICH media. (**b**) For the same cells as in **a**, no influence of ATc was observed on IPTG-induced expression of mScarlet-I. Scale bar 3 μm. (**c**,**d**) Cells were grown at 37 °C in M9A media in 3 independent replicates, and the mean and noise of GFP and mScarlet-I fluorescence intensities (from the average intensities of pixels within cells) were estimated at different IPTG (50, 1250 μM) and ATc (5, 125 nM) concentrations. Translation was inhibited with chloramphenicol for 1 hour prior to preparing microscope samples in order to allow for fluorescent protein maturation. Colored circles indicate the results from independent replicates. For mean expression levels in **c**, black lines indicate the mean GFP and mScarlet-I expression plus or minus 1 standard error of the mean from the 3 replicates. Noise measurements in **d** indicate low noise in both channels in all conditions and replicates.

## Discussion

### Implementing low-noise expression systems

Using modern molecular cloning techniques, it is simple to replace GFPmut2 in pJS101 and pJS102 with genes of interest by PCR and isothermal assembly with near 100% efficiency and a low probability of clones with incorrect sequence. The apparent insensitivity of this circuit to regulatory parameters such as binding affinities for the repressor to DNA and inducer suggests that it can be easily extended to a third, repressor-based expression system. Further, additional ribosome binding sites can be added to the bicistronic operon to express additional components. Notably, we have found that the ATc- and IPTG-inducible systems both function well in MG1655, a strain in which the *lac* operon was deleted, and the *E. coli* TOP10 strain. This host insensitivity could be specific to repressors that are not encoded by the host strain, such as TetR, or repressors that are expressed at very low levels, such as LacI [20].

### Functionality in other organisms

We chose p15a and pSC101 plasmids for these systems because we primarily apply them in *E. coli*, they transform efficiently, and they have been successfully co-transformed in earlier work [6]. However, these are narrow host-range plasmids, and it remains to be seen whether our expression systems will work well in broad host-range plasmids [21], in other organisms, or upon chromosome integration. We expect that the system will be reasonably portable in hosts meeting some basic criteria. First, the P_LtetO-1_ and P_LlacO-1_ promoters are very strong, with sequences close to the *E. coli* σ 70 consensus; this promoter must match promoter sequences recognized in another system. Second, we have created expression systems using a variety of ribosome binding sites with different translation rates [1]; translation rates can be predicted from homology to 16S rRNA and other factors independent of the host [22], and may need to be modified to achieve a desired range of induction. Third, we used the strong, Rho-independent rrnB T1 terminator [23], which should work in a broad range of microbial hosts, but may be less effective in some. Lastly, the addition of an insulating transcriptional repressor ahead of the P_LtetO-1_ and P_LlacO-1_ promoters is likely to reduce sensitivity to transcription upstream of these constructs. We also note that noise for pJS101, with its lower-copy-number pSC101 replicon, is lower than that for pZH509 or pJS102 at similar expression levels (Figure 2b). This suggests that incorporating this construct into the chromosome, where copy number is more tightly regulated, may lead to further noise reduction.

### Possible applications

We expect that the expression plasmids introduced here will be useful for diverse applications in molecular biology. Expression and purification of heteromeric protein complexes could be improved by stoichiometric production of their components, mimicking proportional synthesis in natural systems [24]. Additionally, low-noise expression can improve protein production yields [25]. These systems could also be used in synthetic biology applications where yields can be improved by sequential induction of different components with low cell-to-cell variability. The capacity for low-noise expression at very low expression levels makes them particularly promising for single-molecule imaging experiments or for recombinant expression of low-copy-number components with low cell-to-cell variation to reproduce chromosomal expression levels. We see two major drawbacks to our gene expression system. First, the dynamic range of inducible expression is lower than for systems controlled by constitutively expressed transcriptional repressors [1,7], because some expression must occur at zero inducer concentration before negative feedback kicks in. While the dynamic range could possibly be expanded by increasing repressor binding strength or having a low level of constitutive repressor expression, we have yet to succeed in this. Second, TetR and LacI are expressed at different levels at different induction conditions. This could have off-target effects *(e.g.* from non-specific DNA binding); we have occasionally observed slow growth at very high induction levels (over 200 nM ATc for pZH509), but we have not tested whether this is due to high TetR expression or high GFPmut2 expression.

## Materials and methods

### Strain construction

All plasmids were constructed using isothermal assembly [26] of fragments generated by PCR or double-stranded DNA synthesis (IDT, Coralville) and transformed into Top10 E. coli cells (5-1600-020, IBA Life Sciences, Göttingen). Transformants were screened by colony PCR and verified by sequencing (StabVida, Caparica). Purified plasmids were transformed into *E. coli* strain MG1655 by growing 3 mL of culture in SOB media at 30 °C to an optical density at 600 nm (OD600) of 0.4, washing twice with 1 mL ice-cold water, resuspending in 40 μL water, electroporating 1—10 ng plasmid with the EC1 setting of a Micropulser (Bio-Rad Laboratories, Hercules) and recovering for 1 hour at 37 °C in SOC media before plating on selective LB-agar.

To generate pJS101 with a compatible backbone, plasmid pZH509 [1] was used as a template to amplify the bicistronic regulatory construct including the P_LtetO-1_ promoter [7], GFPmut2 [27], tn10 TetR [28] and rrnB T1 transcription terminator [23]. Using isothermal assembly, this construct was inserted into the pGB2 backbone [5] with the pSC101 origin of replication and spectinomycin resistance to generate plasmid pJS101. Plasmid pJS102 was generated by 3-fragment isothermal assembly. Plasmid pZH509 was used as a template both for the vector backbone and for GFPmut2, with non-homologous extensions added to PCR primers to generate the P_LlacO-1_ promoter [7]. LacI [20] was amplified from *E. coli* MG1655 by colony PCR.

The test strain for mRNA imaging, ZHX99, was constructed similarly to ZHX222 in recent work [29]. In ZHX99, a construct in which a fusion protein of mVenus and Cro is expressed from the bacteriophage λ promoter *P*_*R*_ was integrated into the chromosome to replace the *lac* operon in MG1655 [30]. ZHX99 differs from ZHX222 in three ways. First, the *P*_*R*_ promoter was weakened by site-directed mutagenesis to produce a strain with lower mRNA levels. Second, a very strong ribosome binding site was added (RBS #136 [22]). Lastly, 24 tandem repeats of the recognition sequence for the PP7 coat protein (PP7cp) were inserted between the open reading frame and transcription terminator (amplified by PCR from pDZ251 [17]). The pZH713 plasmid for mRNA detection was constructed by replacing GFPmut2 in pJS102 with a fusion protein of SYFP2 (amplified from a plasmid [31]) and PP7cp (generated after codon optimization by DNA synthesis based on previously reported sequences [32]). Additionally, in pZH713 the PP7cp-SYFP2 fusion protein is translated from the weak ribosome binding site from pZH511 [1]. We note that mVenus expression in ZHX99 is extremely low (undetectable without strong laser excitation) and does not interfere with mRNA detection by localizing up to 48 SYFP2 molecules in a diffraction-limited spot bound to a single mRNA molecule.

To test independent induction of two genes, GFPmut2 in pJS102 was replaced by mScarlet-I (amplified from a plasmid [19]) to make pDG101. Plasmids were co-transformed into MG1655 by electroporation following the above protocol, except with 1 μL each undiluted plasmid (20—40 ng) and selecting on LB-agar plates with both spectinomycin and carbenicillin. Sequence maps are included in **File S2** and plasmids useful for constructing additional two-gene expression systems (pJS101 and pJS102, Table 1) are available from AddGene (deposit numbers 118280 and 118281) and have been verified by whole-plasmid sequencing [33].

**Table 1:**
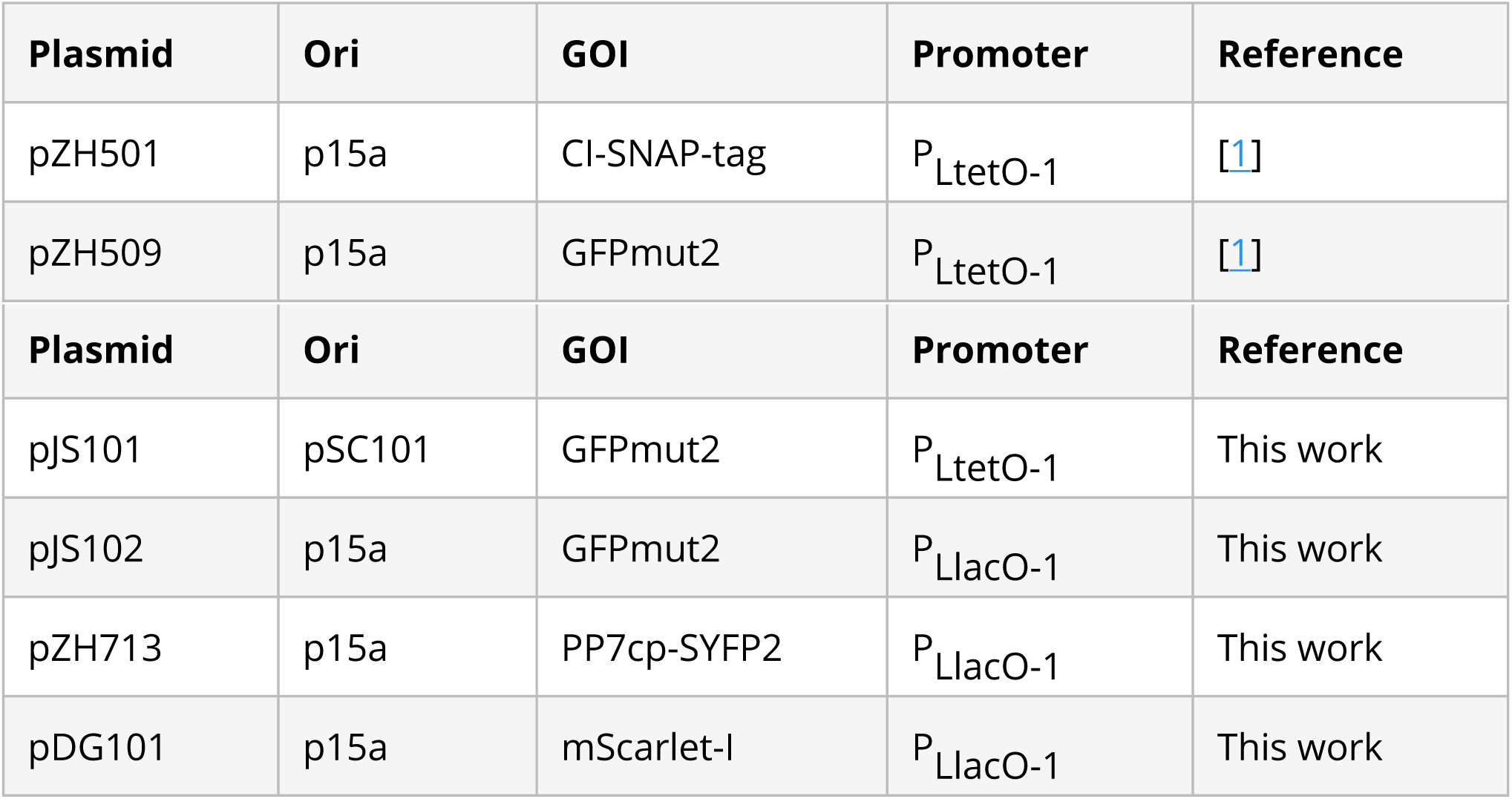
Plasmids used in this study. Ori is origin of replication. GOI is the gene of interest, which in all plasmids is expressed from a bicistronic mRNA with the appropriate repressor (TetR or LacI). Plasmids pZH713 contains a weaker ribosome binding site than other plasmids. Plasmid pJS101 confers spectinomycin resistance and other plasmids confer ampicillin resistance.

### Characterization of GFP expression by flow cytometry

All flow cytometry experiments were repeated 3 times on different days and used plasmids transformed by electroporation into *E. coli* MG1655. Cultures were grown overnight at 30 °C from LB-agar plates supplemented with carbenicillin or spectinomycin (both at 50 *μg/mL*) in 1 mL EZ Rich Defined Medium (M2105, Teknova, Hollister) supplemented with the same antibiotics. Cells were reinoculated 1:400 in 1 mL of the same media supplemented with Isopropyl β-D-1-thiogalactopyranoside (IPTG, at concentrations of 0, 2, 4.5, 10, 22.4, 50, 111.8, 250, 559 and 1250 μM) or anhydrotetrycline (ATc, at concentrations of 1, 5, 25 and 125 nM) as indicated and grown at 30 °C for 4—4.5 hours until reaching an optical density at 600 nm of 0.2—0.3. ATc (Alfa Aesar #J66688) is an analog of tetracycline that binds the tetracycline repressor very strongly, allowing it to be used as an inducer in strains without tetracycline resistance [34]. Next, 10 μL of cells were added to 1 mL of PBS at pH 7.4 and examined by flow cytometry.

Flow cytometry data was collected on an S3e cell sorter (Bio-Rad, Hercules) using a target flow rate of 2,000 counts per second and collecting 30,000 counts for each sample. A 488-nm laser line was used for excitation at its maximum power setting with amplification settings of 450 (forward scattering, FSC), 350 (side scattering, SSC) and 900 (FL1, 525/30 nm). The cell sorter is calibrated daily for a linear response to sample fluorescence intensity. Acquisition was triggered by forward scattering with a threshold of 3. Data was exported as an FCS file and imported into a custom Python script using FlowCal [35]. Following previous methods [1], one third of samples were selected based on proximity to the peak of FSC-AREA and SSC-HEIGHT in a 2D histogram using the density2d method in FlowCal. The FL1-AREA measurements were used to estimate the mean and variance of GFP distributions for all samples. This was done by estimating the probability density functions in bins distributed equally in logarithmic space and fitting by least squares minimization to a gamma function. We found that this method reduced the influence of low-FL1-AREA events that escaped other gating steps, and which had frequencies that varied for different samples and days (Figure S1). In all plots, the mean fluorescence of a strain harboring a similar plasmid, pZH501, that does not encode a fluorescent protein, was subtracted [1].

Noise was calculated as the coefficient of variation squared (*CV*^2^) from the mean, *μ*, and variance, *σ*^2^, as *CV*^2^ = *σ*^2^/μ^2^. We chose *CV*^2^ to facilitate comparison with earlier work [1,36]. We note that gating by FSC-AREA and SSC-HEIGHT to some extent selects for cells near the median cell size, so, ignoring other sources of experimental error, we expect our noise measurements to fall somewhere between the noise in the number of proteins per cell and the noise in the protein concentration. In earlier work identifying the “extrinsic noise limit”, noise was estimated from integrated fluorescence intensities normalized by cell size (proportional to protein concentration) in microscope images [1]. The script for data analysis as well as all raw FCS data is available in **File S2** and utilized modules from SciPy, NumPy, Matplotlib and Pandas.

### Microscopy

All imaging was done on a Leica DMI6000 inverted microscope using illumination from a Leica EL6000 source (at various intensities ensuring minimal photobleaching during acquisition), fluorescence filter cubes (Leica GFP ET, a custom filter set with Semrock filters FF01-561/13, FF02-616/73 and DI02-R561, or the Semrock LF514-B filter set), a 100×/1.46 a-plan apochromat oil immersion objective, Leica Type F immersion oil, and an Evolve 512 EM-CCD camera (Photometrics) using 16-bit EM gain amplification. Images were prepared using Fiji [37], with linear scaling and maintaining minimum and maximum intensity values for all comparable images.

For mRNA imaging, overnight cultures were diluted 1:100 and grown at 30 °C in M9 media supplemented with 1X MEM Amino Acids (M9A) or M9A additionally supplemented with 1% SOB media for 2—4 hours. Supplementation with SOB is used to provide quasi-rich growth conditions with very low fluorescence background and autofluorescence without the expense of commercial rich minimal media. We previously used this growth condition to characterize gene expression noise for different systems [1]. Agarose gel pads (3% BP165-25, Fisher Bio-Reagents) were prepared with M9A with and without supplementation with 1% SOB, and the microscope sample chamber was maintained at 30 °C.

To quantify independent induction from 2 plasmids, cells were grown in M9A at 37 °C. Overnight cultures were diluted 1:100 in M9A supplemented with ATc and IPTG and grown for 3.5 hours. At this point, chloramphenicol was added to a final concentration of 100 μg/mL and cultures were incubated for an additional 1 hour at 37 °C to allow most GFPmut2 and mScarlet-I molecules to mature (maturation times of 5.6 and 25.7 minutes, respectively [38]). Cells were spotted onto agarose gel pads prepared with PBS and imaged at room temperature. For each of 3 replicates and 4 induction conditions, 10 images each were acquired in brightfield, GFPmut2 and mScarlet-I channels. For analysis, all cells in these images were manually segmented using the selection brush tool in Fiji with a width of 10 pixels (163—212 cells per sample). This selection was used to extract the mean green and red intensities (proportional to the concentration of GFPmut2 and mScarlet-I molecules in the cell, respectively). For each image, the mean background intensity was also measured from a large region containing no cells, which was subtracted from each single-cell data point. Mean, variance and *CV^2^* were estimated from this data for each sample following the same fitting protocol used for flow cytometry data. The raw data, single-cell intensities, Fiji macro and Python scripts required to reproduce this analysis are available in **File S2**. Example images were acquired similarly, with growth instead in EZ-Rich media at 30 °C, supplemented with ATc and IPTG.

## Author contributions

JS, SL, DG and ZH designed experiments and performed experiments. JS, SL, DG and ZH performed molecular cloning. ZH, SL and JS analyzed data and wrote the paper. JS, SL, DG and ZH edited and approved the manuscript. ZH supervised the project.

## Acknowledgments

This work was financially supported by: Project LISBOA-01-0145-FEDER-007660 (Microbiologia Molecular, Estrutural e Celular) funded by FEDER funds through COMPETE2020—Programa Operacional Competitividade e Internacionalização (POCI), by national funds through FCT—Fundação para a Ciência e a Tecnologia, and through a joint research agreement with the Okinawa Institute of Science and Technology (OIST). This manuscript was composed and edited using Manubot [39].

## Supplementary Material

**Figure S1:**
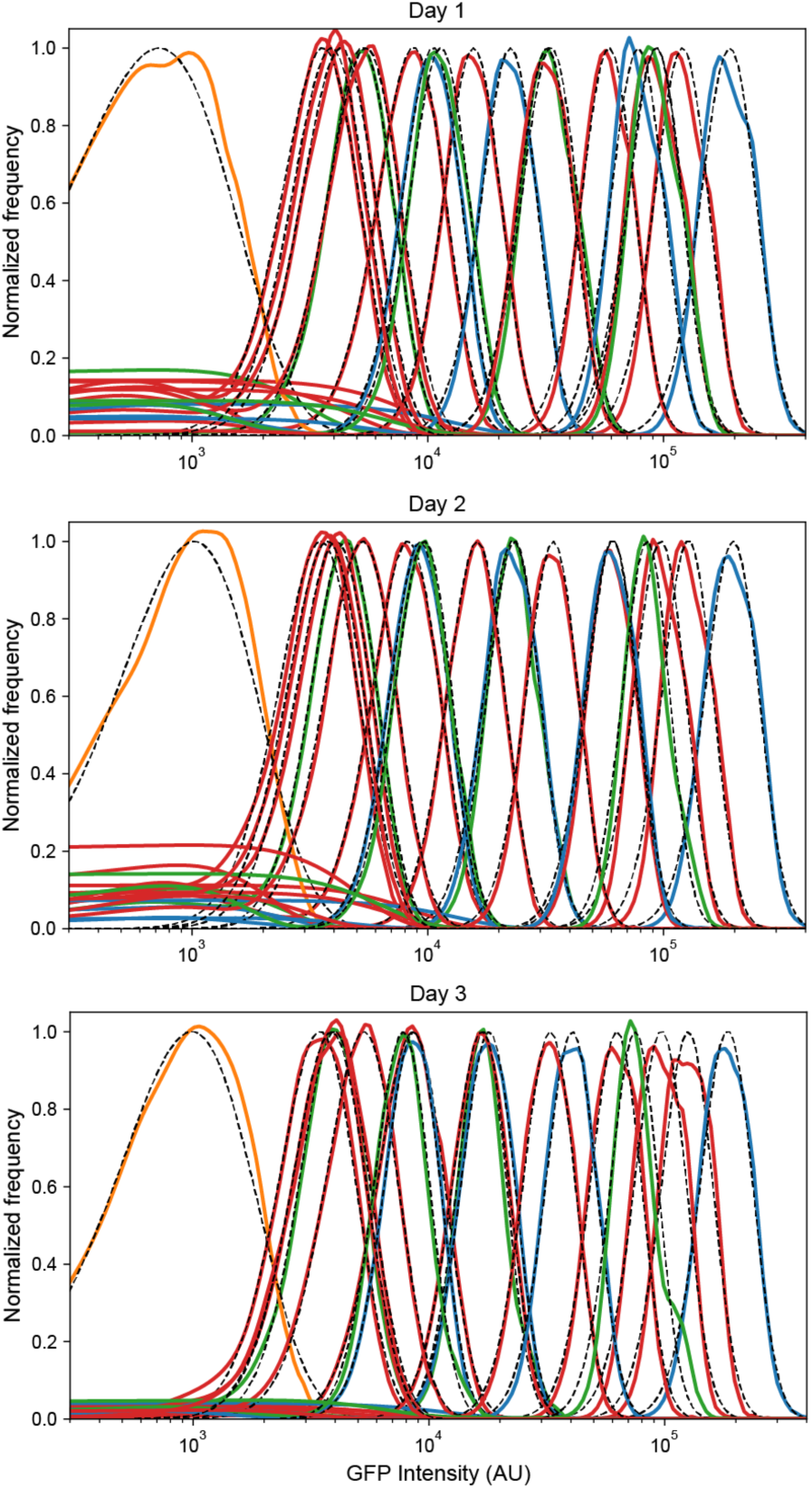
Reproducibility of low-noise expression in 3 independent experiments. Probability densities for each flow cytometry sample were calculated by kernel density estimates for the negative control plasmid pZH501 (orange), ZH509 (blue), pJS101 (green), and pJS102 (red) with fluorescence levels monotonically increasing with concentration of ATc (1, 5, 25, 125 nM) or IPTG (0, 2, 4.5, 10, 22.5, 50, 111.8, 250, 559, 1250). Distributions were fit by least squares regression to a gamma function (black dashed lines) to estimate sample mean and variance while minimizing the influence of non-fluorescent background events, which varied in frequency for different days and samples. All plots are normalized by the maximum value of the fit gamma distribution.

**Supplementary File S2**: Raw flow cytometry data, Python scripts required to reproduce Figures 1, 2, and 4, DNA sequences, and explanatory text files are available as a compressed archive at Zenodo [40].

